# The Effect of Ethanolic Extract of *Acacia Seyal* Bark on Induced Diabetic rats

**DOI:** 10.1101/2022.01.21.476925

**Authors:** LM Elamin Elhasan, Basmat Elhkotam, Tomader Salah Abdelgadir, Smaher Greeb Allah Ibraheim, Omar Musa Izz Eldin Othman

## Abstract

**Background:** Diabetes mellitus Type 2 is a chronic metabolic disorder characterized by insulin insensitivity that leads to a decrease in glucose transport into the other cells. Many drugs are being developed, while no cure is available regarding to this disease although, there are limitations due to high cost and certain side effects. The Traditional Medicines are preferred due to lesser side effects and low cost. Acacia species have wide traditional medicinal using as anti-diabetic, antimicrobial, and anti-inflammatory activities.

**Objective:** The study aimed to examine the effect of ethanolic extract of *Acacia Seyal* bark in induced diabetic rats.

**Materials and methods:** *Acacia Seyal* bark was extracted in ethanol 80%. The Ethanolic extract was analyzed for phytochemicals screening tests using standard methods. To investigate the effect of the extract thirty induced diabetic albino rats introduced by injection of glucose 2g\kg were divided to 5 groups equally; Group 1 was treated with 10mg\kg of glibenclamide, group 2 left as control treated with distilled water 10mg\kg, Group 3 was treated with 200mg of plant extract, Group 4 was treated with 400mg of plant extract, and Group 5 was treated with 800mg of plant extract. The Glucose Tolerant Test were done after 1 hour, 2 hour and 4 hour to determine blood glucose level of rats. Estimation of in vitro glucose uptake by rat diaphragm experiment was done to evaluate the glucose utilization capacity of extract.

**Results:** The effect of different concentration of *ethanolic* extract of *Acacia Seyal* bark on the blood glucose level of diabetic induced rats is significantly different (0.05 p 0.01 <p). The phytochemical screening of extract indicated the presence of Saponin, Tannins Steroids, Triterpens, and Anthraquinone, While the absence of flavonoids. In vitro glucose consumption by diaphragm study the level of utilized glucose from the media is 69.4% in the presence of extract compared to control group.

**Conclusion:** Study concluded that the ethanolic bark extract of *A. seyal* showed significant anti-diabetic activity. This results might have a great potential for translation to humans and the obtained data might set the stage for clinical trials investigating the effects of anti-diabetic effect in patients with diabetes mellitus type2. In the best of our knowledge, this is the first study that has conducted to study the anti-diabetics effect of ethanolic extract of *Acacia Seyal* bark in diabetic rats.

## 1 Introduction

The incidence of Diabetes mellitus Type 2 (DM) has been increasing over the world, It is estimated that 439 million people would have type 2 DM by the year 2030 [1, 2].

Diabetes mellitus Type 2 (DM) is a chronic metabolic disorder characterized by insulin insensitivity as a result of insulin resistance, declining insulin production, and eventual pancreatic beta-cell failure [3, 4]. that leads to a decrease in glucose transport into the liver, muscle cells, and other cells [5]. The pathophysiological changes of DM type 2 occur in several years before its detection. So early detection and treatment of disease can suppress disease’s progression, complications, and prevent beta-cell dysfunction [6].Early screening and diagnosis of DM type 2 are prevent of disease for 98%, the result that a positive screen is equivalent to a diagnosis of pre-diabetes or DM[7], Although about 25% of patients with type 2 DM already have microvascular complications at the time of diagnosis suggesting that they have had the disease for more than 5 years [8]. Type 2 DM can be prevented with lifestyle modification; exercise and diet control and community awareness about the disease.[2].

Many drugs are being developed, while no cure is available regarding to this disease [2] Although, there are limitations due to high cost and certain side effects such as development of hypoglycemia, weight gain, gastrointestinal disturbances, liver toxicity [9], efforts are on to find suitable anti-diabetic therapy.

The Traditional Medicines from medicinal plants which used by about 60% of the world’s population, are preferred due to lesser side effects, availability, and low cost. A list of medicinal plants have anti-diabetic effects[10]. In example Acacia species have medicinal benefits as anti-diabetes[11, 12] and anti-microbial[13, 14].

*Acacia seyal* belongs to the family of Fabaceae (Mimosoideae). It is a small to medium-sized tree and up to 17 m tall and 60 cm in diameter, it is widespread in the semi-arid zone of tropical Africa and the Red Sea and from the Nile valley south to Zambia[12, 13]. In Sudan, *A. seyal*, locally known as Talha[14], It is one of the most important medicinal plants used in traditional, or alternative, medicine, fodder sources for livestock in Sudan[15], and source of gum Arabic. Gum Arabic (GA) is a mixture of polysaccharides and glycoproteins secreted from *Acacia senegal, Acacia seyal*, and *Acacia nilotica* trees’ stems[16] Pharmacologically, GA has been confirmed to have several therapeutic actions, such as being hypoglycemic, antidiabetic, antioxidant, immunomodulatory, antiulcer, protect against hepatic, renal, and cardiac complications in diabetic and chronic renal failure patients[17].

## 2- Materials and methods

### 2-1 Plant collection

The bark of *Acacia Seyal* was collected from Kordufan in western region of Sudan. The bark was washed under tap water and completely shade dried at room temperature. Then ground into fine powder by mortar and pestle. The powders were packed in clean plastic bag.

### 2-2 Preparation of Ethanolic extract

Extraction was carried out according to method described by Sukhdev [18] 100 g was coarsely powdered using mortar and pestle. Coarsely sample was extracted with ethanol 80% using Soxhlet extractor apparatus. Solvents were evaporated under reduced pressure using rotary evaporator apparatus. Finally extracts allowed to air in Petri dish till complete dryness.

### 2-3 phytochemical screening tests

Phytochemical screening for the active constituents was carried out on Methanolic extract using the methods described by (Martinez & Valencia (1999), Sofowora (1993), Harborne (1984) and Wall et al (1952)) with many few modifications [19-22].

#### 2-3-1 Identification of tannins

0.5 g of the extract washed three times with petroleum ether, dissolved in 10 ml hot saline solution and divided in two tests tubes. To one tube 2-3 drops of ferric chloride added and to the other one 2 – 3 drops of gelatin salts reagent added. The occurrence of a blackish blue colour in the first test tube and turbidity in the second one denotes the presence of tannins.

#### 2-3-2 Test of sterols and triterpenes

0.5 g of the extract washed three times with petroleum ether and dissolved in 10 of chloroform. To 5 ml of the solution, 0.5 ml acetic anhydride was added and then 3 drops of conc. Sulphuric acid at the bottom of the test tube. At the contact zone of the two liquids a the gradual appearance of green, blue pink to purple color was taken as an evidence of the presence of sterols (green to blue) and or triterpenes (pink to purple) in the sample.

#### 2-3-3 Tests for Flavonoids

0.5 g of the extract was washed three times with petroleum ether, dissolved in 30 ml of 80% ethanol. The filtrate was used for following tests: -

A/ to 3 ml of the filtrate in a test tube 1ml of 1% potassium hydroxide solution in methanol was added. Appearance of a yellow color indicated the presence of Flavonoids. Flavones or and chalcone.

B/ to 2 ml of the filtrate 0.5 ml of 10 % lead acetate was added. Appearance of creamy turbidity was taken as an evidence of Flavonoids.

#### 2-3-4 Test for Saponins

0.3 g of the extract was placed in a clean test tube. 10 ml of distilled water was added, the tube stoppered and vigorously shaken for about 30 seconds. The tube was then allowed to stand and observed for the formation of foam, which persisted for least an hour, was taken as evidence for presence of saponins.

#### 2-3-5 Test for Anthraquinone glycoside

0.5 g of the extract was boiled with 10 ml of 0.5N KOH containing 1ml of 3% hydrogen peroxide solution. The mixture was extracted by shaking with 10 ml of benzene. 5ml of the benzene solution was shacked with 3ml of 10% ammonium hydroxide solution and the two layers were allowed to separate. The presence of Anthraquinone was indicated if the alkaline layer was found to have assumed pink or red color.

### 2-4 Laboratory animals

Thirty albino mice (80-120g) were housed in polypropylene cages inside a well-ventilated room at 20 ± 4 °C and 12-h periods of light and dark, with free access to clean water and commercial mice food. All mice were fasted for 18 h before experimental induction of diabetes. At the end of experimental period, animals were sacrificed.

### 2-5 Induction of hyperglycemia

Diabetes was induced experimentally by administration of 50% glucose. The animals were fasted for 18 hours, but allowed free access to water. A dose of 2 g/kg body weight glucose was administered by intraperitoneal injection, blood glucose level was determined by use of colorimeter.

### 2-6 Experimental design

The rats were divided in to five groups, each group contain 6 rats. Group 1 was treated with 10mg\kg of glibenclamide, group 2 left as control treated with distilled water 10mg\kg. Group 3 was treated with 200mg of plant extract. Group 4 was treated with 400mg of plant extract. Group 5 was treated with 800mg of plant extract.

### 2-7 Blood sampling

Blood samples were collected from the rats using orbital techniques to determined blood glucose level. Blood samples were collected from the retro bulbar plexus of the median canthus of the eye of the rats. A micro capillary tube was carefully inserted into the medial canthus of the eye to puncture the retro bulbar plexus and thus enable out flow of about 2 ml of blood into a blood glucose test tube. Then, test tubes were centrifuged at 3,000 rev per min using a table centrifuge. The Glucose Tolerance Test. was done at the start of the experiment and after 1, 2, and 4 hours [23].

### 2-8 Estimation of in vitro glucose uptake by rat diaphragm

Glucose uptake by rat diaphragm in vitro was estimated by a standard method. Two rats were weighing between 170 to 210 g were used in this study. The rats were anesthetized and dissected and cut into two equal halves. The diaphragms were dissected out quickly with minimal trauma and divided into2 halves. The diaphragms were rinsed in cold tyrode solution Tyrode (NaCl (8 gm/L), KCl (0.20 gm/L), CaCl2 (0.20 gm/L), MgCl2 (0.10 gm/L), NaH2PO4 (0.05 gm/L), NaHCO3 (1 gm/L), Glucose (1 gm/L) having pH 6.5) without glucose to remove any blood clots and were placed in small culture tubes containing 2ml tyrode solution. Four sets containing of incubating tubes were prepared as followed; Tube I which called PRO contained of 2ml of tyrode solution and 1 ml of insulin (0.25IU/ml), tube II which present the control contained of 2ml of tyrode solution, tube III contained of 2 ml of tyrode solution and 1 ml of extract, tube IV contained of 2ml of tyrode solution, 1 ml of extract, and 1 ml of Insulin. The volumes of all tubes were made up to 4 ml with distilled water to match the volume of the tube IV.

Each diaphragms were placed in incubating tubes and incubated for 30 min at 37°C in an atmosphere of 95% oxygen, and glucose 2% with shaking at 140 cycles/min. Glucose uptake per gram of tissue was calculated as the difference between the initial and final glucose content in the incubated medium.

### 2.9 Data analysis

Differences between the means of the various groups of in the efficacy study was done using ANOVA statistical test, while the differences between the means of the two groups used in the effect of different concentration of *Acacia Seyal* bark ethanolic extract were done using students t-test. Level of significance for all the analysis was set at P<0.05.

## 3- Results

### 3-1 Yield Percentage of Ethanolic Extraction of *Acacia Seyal* Bark

The yield percentage of Ethanolic Extraction of *Acacia Seyal* bark were calculated as followed: Weight of extract obtained / weight of plant sample X100, that shown in table I.

**Table I:** The Yield Percentage of Ethanol Extraction of *Acacia Seyal* bark.

| Sample | Weight of sample in gm | Weight of extract | Yield % |
| --- | --- | --- | --- |
| *Acacia Seyal* | 50 g | 5.242 | 10.5 % |

### 3-2 phytochemical screening test of Ethanolic Extract of *Acacia Seyal* bark

Qualitative screening of Ethanolic extract of *Acacia seyal i*ndicated the presence of sponin, tannins, steroids, triterpens and anthraquinone and the absence of flavonoide. That shown in table II.

**Table II:**
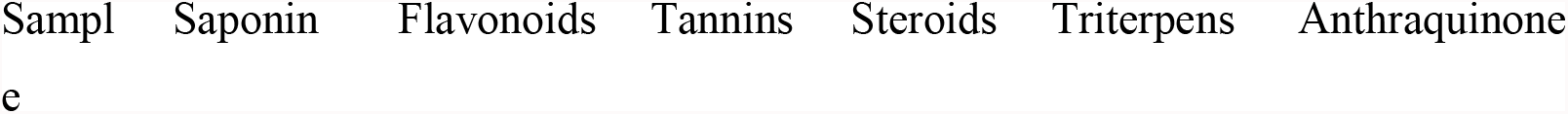
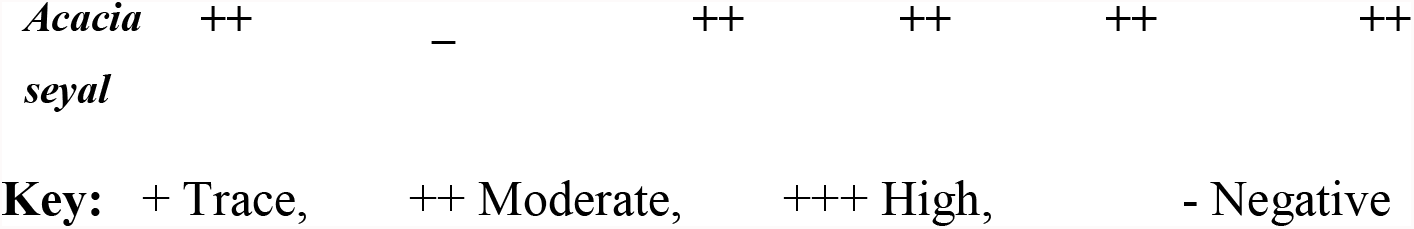
Phytochemical screening test of Ethanolic extract of *Acacia Seyal* bark.

### 3-3 The effect of different concentration of Ethanolic extract of *Acacia Seyal* bark

The blood glucose level of diabetic rats is significantly different (0.05 p 0.01 <p) after 1 hour, 2 hours, and 4 hours from the initiation of the experiment was determine by Glucose Tolerant Test using calorimeter, that shown in table III

**Table III:** Effects of orally administered different concentration of Ethanolic extract of *Acacia seyal* in diabetic rats and the level of blood glucose, using t test.

|  | **1hr** | **2hr** | **4hr** |
| --- | --- | --- | --- |
| Control/ | 121.3 | 113.4 | 181.5 |
| glibenclamide | 132.3 | 140.3 | 139.7 |
| 200 mg/kgbw | 83.7 | 100.2 | 114 |
| 400mg/kgbw | 85.7 | 96 | 123 |
| 800mg/kgbw | 70 | 81.8 | 96.3 |
| L.S.D | 30.7 | 24.3 | 37.4 |

Previous results are expressed as Means ± SD for five animals per group. Values followed by the same superscript are statistically different (P ≤ 0.05; Analysed by ANOVA test that shown in table IV.

**Table IV:**
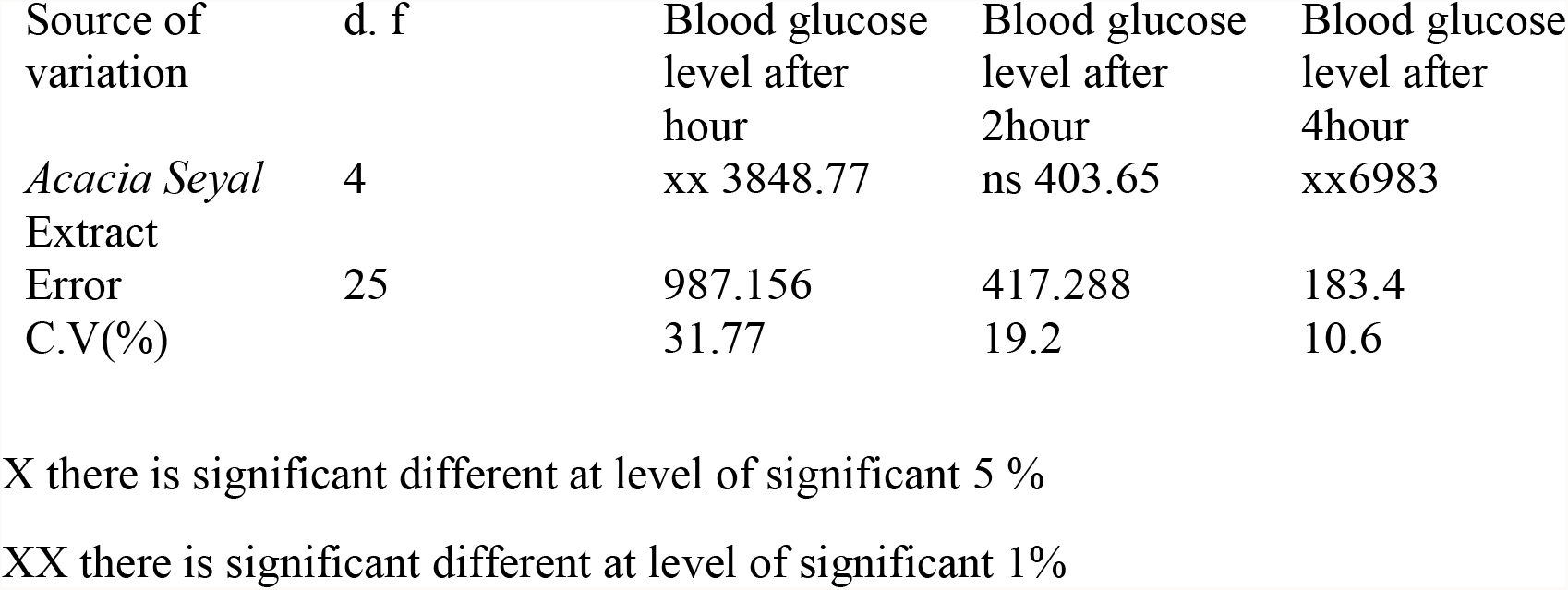
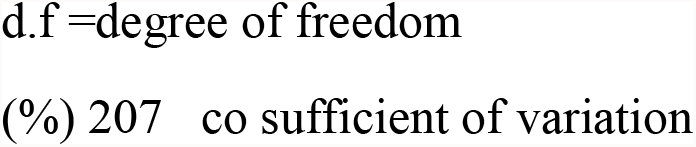
Table IV: The value of Means ± SD of the effect of different concentration of Ethanolic extract of *Acacia Seyal* analyzed by ANOVA test.

### 3-4 Estimation of in vitro glucose uptake by rat diaphragm

To estimate the Glucose uptake per gram of tissue was calculated as the difference between the initial and final glucose content in the incubated medium. The glucose content of the incubated medium was measured by colorimeter using GOD-POD method.

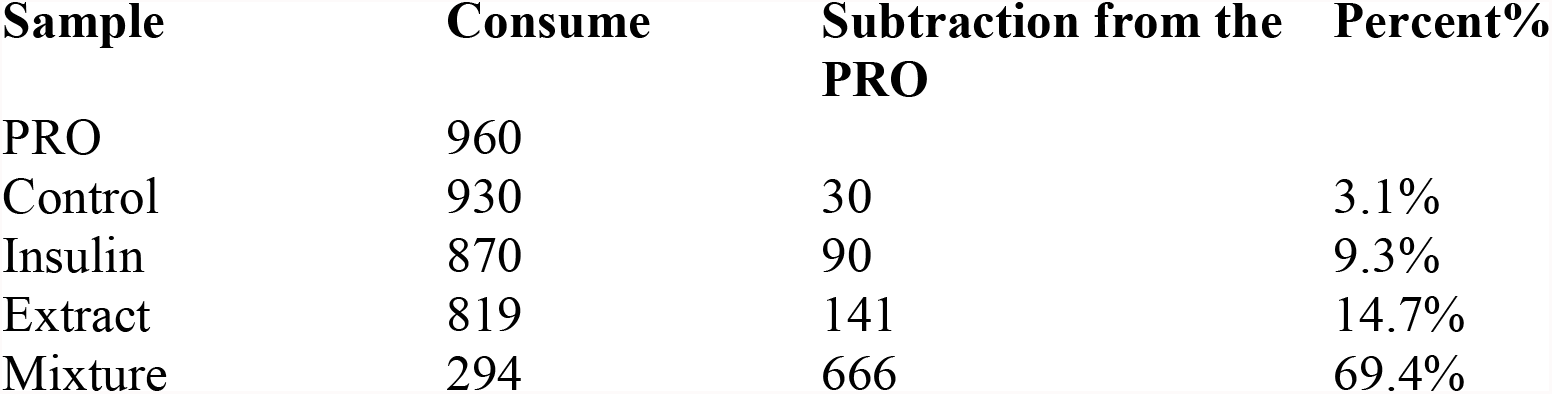

## 4- Discussion

This is the first study on the anti-diabetic effect of ethanolic extract of *Acacia Seyal* bark in induced diabetics’ rats caused by injection of glucose 2g\kg. The experimental rats weighting (80-120g) were used in this study. The rats were divided in to five groups, each group contain 6 rats. Group 1 was treated with 10mg\kg of glibenclamide, group 2 left as control treated with distilled water 10mg\kg. Group 3 was treated with 200mg of ethanolic extract of *Acacia Seyal* bark. Group 4 was treated with 400mg of plant extract. Group 5 was treated with 800mg of plant extract. The standard group, the group which treated with glibenclamide cause hypoglycemia reduced the blood glucose level of diabetic rats to (140.3-139.7) at the time interval 2,4h respectively. In the group which treated with plant extract at dose 800mg\kg reduced blood glucose level of diabetic rats to 81.3-96.3 at the time interval 2,4h respectively. In comparison with the group which treated the plant extract at dose 400mg\kg showed mordent decrease in the blood glucose level of diabetic rats to 96-123at the time interval 2,4h respectively. the group which treated the plant extract at dose 200mg\kg given least in blood glucose level of diabetic rats to 100.2-114 at the time interval 2,4h respectively, this compatible with the other studies which used extract from different species of Acacia; Acacia farnesiana (L.) willd water extract at a dose of 50 mg/kg showed significant glucose lowering activity at 30 and 90 min after glucose loading in normal fed rats [24]. Study by Rahmatullah M[25] which used *Acacia catechu* prepared from boiling the wood in water the different doses of extract was administered. The study concludes significant oral hypoglycemic activity on glucose-loaded mice at all doses of the extracts tested but shown less effect than that of a standard drug, glibenclamide (10 mg/kg body weight). Another study used mixture of hops rho iso-alpha acids-rich extract and acacia proanthocy-anidins-rich extract prove that Mice fed by the mixture extract (100 mg/kg) for 7 days decrease in non-fasting glucose 22%. and 19% decrease in insulin that was comparable to 0.5 mg/kg rosiglitazone and better than 100 mg/kg metformin [26].

To estimate how the Glucose consumed from the media in vitro by isolating diaphragm from albino rats. Each diaphragms were divided into2 halves and placed in incubating tubes. Four sets containing of incubating tubes were prepared as followed; Tube I which called PRO contained of 2ml of tyrode solution and 1 ml of insulin (0.25IU/ml), tube II which present the control contained of 2ml of tyrode solution, tube III contained of 2 ml of tyrode solution and 1 ml of extract, tube IV contained of 2ml of tyrode solution, 1 ml of extract, and 1 ml of Insulin. In the first tube which called (pro) the level of utilized glucose from the media is 3.1%. In the second group which called (control) the level of utilized glucose from the media is 9.3%. In third tube which called (extract) the level of utilized glucose from the media is 14%. In fourth tube which called (mixture) the level of utilized of glucose from the media is 69.4% significantly increased glucose uptake by isolated rat diaphragm as compared to control, that agree with previous study which used Acacia farnesiana (L.) extract the isolated hemidiaphragm, glucose uptake was increased by treatment at 40 μg/ml of extract [24].

The phytochemical screening of ethanolic extracts indicated the presence of Saponin, Tannins Steroids, Triterpens, and Anthraquinone, While the absence of flavonoids. Tannin, or tannic acid, activates glucose transfer and prevents lipolysis [27]. The results of study by Elmi, A [28] suggest that *A. seyal* bark could be a potential source of active natural compounds like flavonoids, steroids, triterpenes, and tannins as catechin and gallocatechin-based oligomers. Gums are consist of (heteropolysaccharides) and condensed tannins (flavan-3-ol derivatives) are the most commonly reported constituents in Acacia. Pharmacological studies prove the antioxidant, analgesic, antihypertensive, antidiabetic, anti-Alzheimer, anti-microbial [29, 30] and antimalarial effects of Acacia extracts. *Acacia species* constitutes of certain secondary metabolites including phenols, alkaloids, and terpenoids, some with useful biological activities, have been reported in acacias. Very few species have been investigated for their phytochemical composition and biological activities [31].

In the best of our knowledge, no study has ever evaluated the hypoglycemic effects of *A. seyal* Bark in type II DM rats. Further scientific studies to isolate of bio-active constituents are highly recommended.

## In conclusion

Study concluded that the ethanolic extract of *A. seyal* bark showed significant anti-diabetic effect. In this study, In vitro glucose consumption by diaphragm the level of utilized glucose significantly higher in the presence of the extract.

Further, qualitative and quantitative phytochemical screening of *Acacia Seyal* bark to determine the active constitutes are recommended.

This results might have a great potential for translation to humans and the obtained data might set the stage for clinical trials investigating the effects of anti-diabetic effect in patients with diabetes mellitus type2. In the best of our knowledge, this is the first study that has conducted to study the anti-diabetics effect of ethanolic extract of *Acacia Seyal* bark in introduced diabetic rats.

